# Atlas-scale Single-cell DNA Methylation Profiling with sciMETv3

**DOI:** 10.1101/2024.08.29.610369

**Authors:** Ruth V. Nichols, Lauren E. Rylaarsdam, Brendan L. O’Connell, Zohar Shipony, Nika Iremadze, Sonia N. Acharya, Andrew C. Adey

## Abstract

Single-cell methods to assess DNA methylation have not yet achieved the same level of cell throughput compared to other modalities. Here, we describe sciMETv3, a combinatorial indexing-based technique that builds on our prior technology, sciMETv2. SciMETv3 achieves nearly a 100-fold improvement in cell throughput by increasing the index space while simultaneously reducing hands-on time and total costs per experiment. To reduce the sequencing burden of the assay, we demonstrate compatibility of sciMETv3 with capture techniques that enrich for regulatory regions, as well as the ability to leverage enzymatic conversion which can yield higher library diversity. We showcase the throughput of sciMETv3 by producing a >140k cell library from human middle frontal gyrus split across four multiplexed individuals using both Illumina and Ultima sequencing instrumentation. This library was prepared over two days by one individual and required no expensive equipment (e.g. a flow sorter, as required by sciMETv2). The same experiment produced an estimated 650k additional cells that were not sequenced, representing the power of sciMETv3 to meet the throughput needs of the most demanding atlas-scale projects. Finally, we demonstrate the compatibility of sciMETv3 with multimodal assays by introducing sciMET+ATAC, which will enable high- throughput exploration of the interplay between two layers of epigenetic regulation within the same cell, as well as the ability to directly integrate single-cell methylation datasets with existing single-cell ATAC-seq.

**Highlights:**

- Atlas-scale production of single-cell DNA methylation libraries in a single experiment
- Protocols and evaluation using both Illumina and Ultima Genomics sequencing platforms
- Compatibility of sciMETv3 with capture techniques to reduce sequencing burden
- Compatibility of sciMETv3 with enzymatic conversion methods
- Generation of an integrated >140,000 cell dataset from human middle frontal gyrus across four individuals
- Ability to profile both ATAC and genome-wide DNA methylation from the same cells and integration with datasets from each modality
- A novel implementation of the s3-ATAC technology that leverages a nanowell chip for increased throughput

**Motivation:** DNA methylation forms a basal layer of epigenomic regulatory control, shaping the genomic permissiveness of mammalian cells during lineage specification and development. Aberrant DNA methylation has been associated with myriad health conditions ranging from developmental disorders to cancer. The high cell type specificity necessitates analysis at the single-cell level, much like transcription or other epigenomic properties. However, robust and cost-effective techniques to produce atlas-scale datasets have not been realized for DNA methylation. Here, we directly meet this need by introducing sciMETv3, a high-throughput protocol capable of producing hundreds of thousands of single-cell DNA methylation profiles in a single experiment.

## Introduction

Mammalian DNA methylation takes the form of a methyl group covalently added to the 5-carbon of cytosine residues in the genome and forms the most basal layer of gene regulatory control, with distinct programs that shape the permissible genomic landscape during development. Historically, DNA methylation has been profiled using ‘conversion’-based approaches, which leverage chemical or enzymatic processes to convert non-methylated cytosines to uracil. Converted bases are then sequenced as thymine, whereas methylated cytosines are protected from this process. The complexity of conversion protocols makes single-cell approaches particularly challenging, with most methods requiring the deposition and processing of individual cells into their own reaction compartments for conversion and then initial processing steps^1–5^. We previously developed techniques to increase the cell throughput for profiling DNA methylation, sciMET^6^ and sciMETv2 ^7^, which leverage single-cell combinatorial indexing to pre-index cells prior to conversion and the final stages of library preparation. This workflow enables the production of thousands of single-cell methylation libraries to be produced by a single individual and amortizes reagent costs over many pre-indexed cells, substantially reducing costs per cell. We also demonstrated the ability to perform target capture on regulatory loci with high levels of expected cell type specific methylation variability (sciMET-cap) which reduces the number of sequencing reads required per cell to achieve cell type identification and robust cell type clustering^8^.

The sciMETv2 technology can achieve a modest scale of throughput, with typical experiments producing between 5 and 20 thousand single-cell profiles. This capacity is suitable for many applications; however, to achieve the higher end of that range multiple plates of indexed tagmentation must be performed which can be cumbersome and expensive. Here, we directly address these remaining challenges by developing sciMETv3, which leverages an additional tier of cell barcoding to increase throughput by orders of magnitude. Final cell count is flexible and spans three orders of magnitude from ∼1,000 to up to 10 million in increments of ∼1,000 cells. This technology requires comparable hands-on time to sciMETv2 and produces an identical molecular structure, allowing for capture techniques to be carried out. We further demonstrate the ability to perform enzymatic conversion, as well as a modified workflow to enable libraries to be sequenced on the Ultima Genomics platform. We then combine datasets sequenced by both platforms to produce >140,000 cells from the middle frontal gyrus across four healthy human donors. Finally, we demonstrate a variant of the technology that employs two rounds of indexed tagmentation followed by sciMETv3 processing to capture ATAC plus genome-wide DNA methylation profiles from the same cells in high throughput.

## Results

### sciMETv3 design

To achieve increased throughput for the sciMET platform, we devised a strategy to incorporate an additional round of indexing post tagmentation and prior to distribution into the final PCR-indexed wells (Fig. 1A). This approach was based on a ligation workflow similar to that which was achieved for sci-ATAC-seq3^9^. Ligation adapters were designed to directly append to the transposase adapter sequence, completing the 5’ half of the Illumina read 2 sequencing primer. These adapters also append a well-specific barcode and terminate with the Illumina flowcell primer sequence at the 5’ end. The final ligation product results in the same final molecular structure that is produced during PCR for the sciMETv2 workflow, retaining compatibility with downstream capture methods (Fig. 1B,C). The ligation adapters must survive bisulfite conversion and were therefore fully methylated at all cytosine positions. As an initial assessment, we leveraged a set of 96 indexed primers, effectively increasing the throughput of the sciMETv2 platform by 96-fold. The workflow was carried out on four human brain specimens (cortex, BA 46; 90% of nuclei) and a mouse brain specimen (whole brain, C57BL/6; 10% of nuclei), allowing us to estimate our cell doublet rate while providing enough human cells for an initial analysis (Fig. 1D). We then processed a single final PCR well out of a total of 8 that were diluted, which produced 293 passing cell profiles with a mean unique read count of 354,763 and a mean coverage of 2.73 million total cytosines covered per cell. Of these, 269 were human and 24 were mouse, with zero cells identified as doublets, establishing a maximum doublet bound of 3.4% when factoring in the 10-fold skewing toward human cells (Fig. 1E). We next assessed crosstalk by measuring the percentage of cross-species aligned reads, also adjusting for the skewed species mixture, resulting in a maximum of 1.2%. Human cells were taken through windowing and clustering. Leveraging both mCG and mCH contexts produced two neuronal and three glial clusters which were annotated based on global CH methylation levels (Fig. 1F).

**Figure 1.**
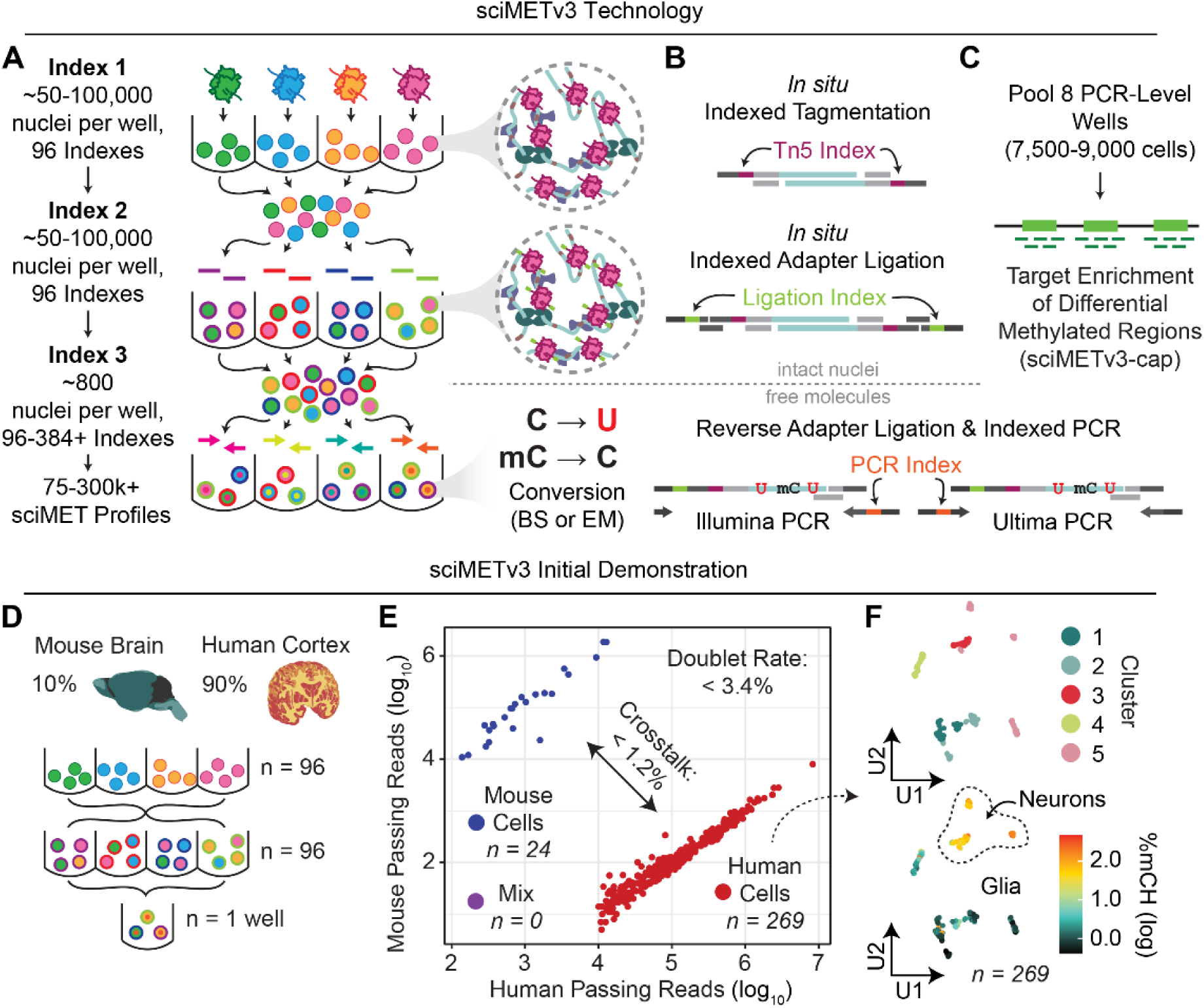
sciMETv3 technology development. **A)** Indexing and pooling schematic for sciMETv3. **B)** Molecular schematic. **C)** Strategy for sciMET-cap enrichment strategy. **D)** Experimental design schematic for initial sciMETv3 development. **E)** Assessment of doublet rates and cell-cell crosstalk from human and mouse cells. **F)** UMAP of human cells from the initial experiment reveals clear clusters (top) with expected mCH patterns for glia and neurons (bottom).

### sciMETv3 is compatible with enzymatic conversion methods as well as target capture

We next assessed the full workflow and platform versatility of sciMETv3 by carrying out a preparation on a human brain specimen (cortex, BA 46). We leveraged 96 tagmentation and ligation indexes and distributed a target of 750 pre-indexed nuclei into each well of a final plate (Fig. 2A). Unlike sciMETv2, the greater number of nuclei within each final well allows for dilution to be deployed as opposed to flow sorting, reducing the overall time of the experiment and eliminating the need for flow cytometry instrumentation. Dilution has been developed for a commercialized version of a combinatorial indexing based single-cell methylation workflow; however, the increased nuclei count of sciMETv3 provides greater robustness at this stage. Eight wells were taken through bisulfite conversion, reverse adapter ligation and PCR. All eight wells (estimated cell n = 6,000) were taken through the capture workflow followed by sequencing, producing 5,805 QC-passing single-cell DNA methylation profiles with a comparable target fold enrichment to sciMET-cap (6.2-fold versus 7 to 10-fold^8^).

**Figure 2.**
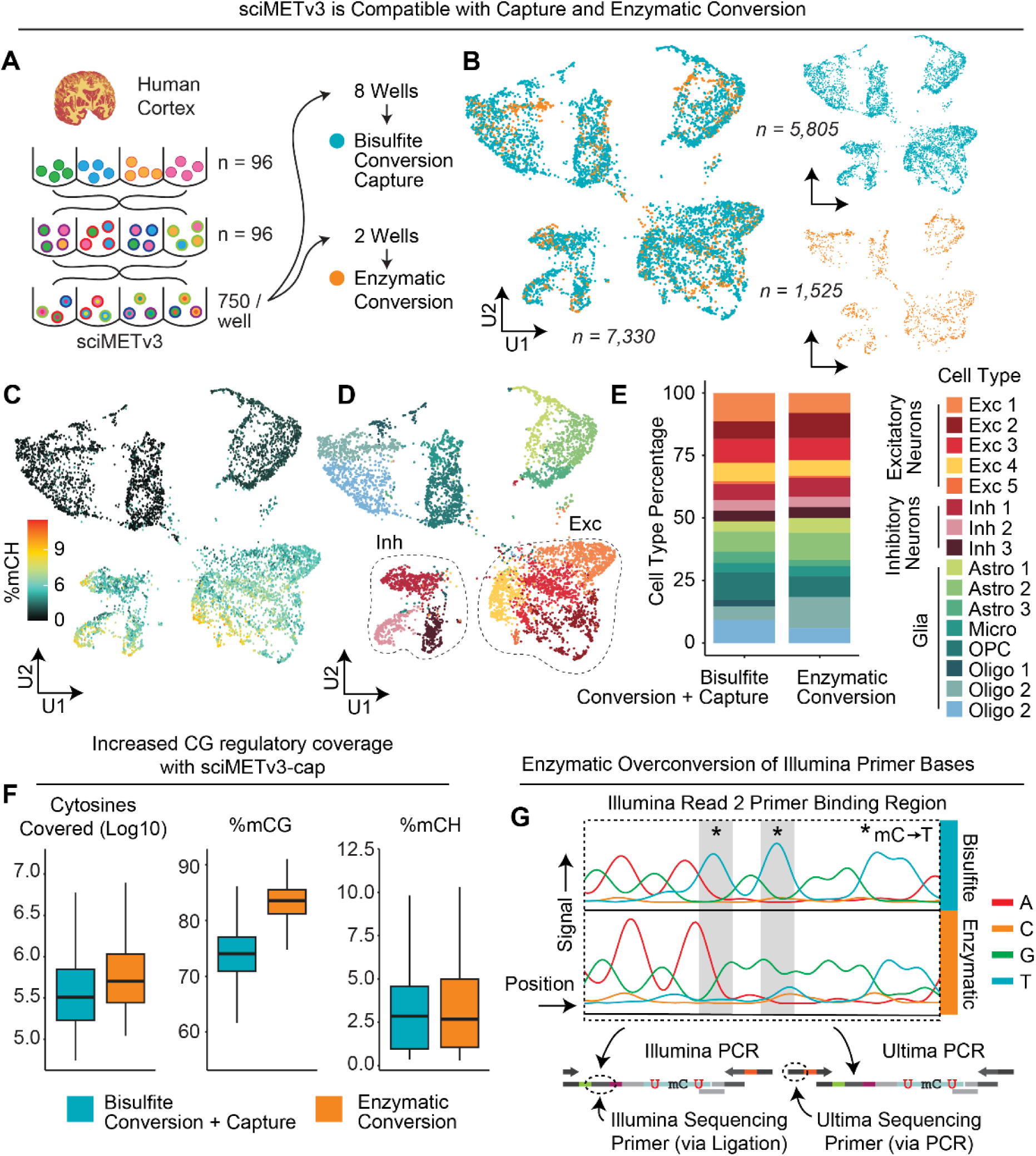
sciMETv3 is compatible with capture methods and enzymatic conversion. **A)** Experimental design schematic. **B)** UMAP of cells combined from both sciMET-cap using bisulfite conversion and non-captured enzymatic conversion preparations. **C)** mCH levels show expected patterns for neurons and glial cell populations. **D)** Identified clusters with inhibitory and excitatory neuron clusters highlighted. **E)** Cluster proportions are comparable between bisulfite + capture and enzymatic non-captured conditions. **F)** Global methylation patterns show expected trend with cells taken through capture exhibiting lower mCG levels due to the enrichment at regulatory loci with no impact on mCH levels. **G)** Sanger sequencing traces of enzymatic converted libraries show over-conversion of key bases present in the read 2 / index read 1 Illumina sequencing primer region that is appended during adapter ligation.

The increased nuclei count per conversion well for sciMETv3 over sciMETv2 (96-fold greater) brings the total input within the recommended range for enzymatic conversion methods without the need for ultra- low-input modifications. Enzymatic conversion methods have been shown to offer improved yields over the harsh chemical processes of bisulfite conversion^10^ and have been demonstrated previously in the context of other sciMET-like protocols^11^. Two wells of the final plate (estimated cell n = 1,500) were taken through enzymatic conversion followed by reverse adapter incorporation and PCR. Sequencing produced 1,525 QC- passing single-cell methylomes. As anticipated, the insert size of library fragments from the enzymatic conversion library were greater than that of bisulfite methods (mean = 163 ± 121 vs 78 ± 69 bp for enzymatic and bisulfite, respectively; 2.1-fold increase).

We next aggregated cell profiles from both the bisulfite-converted sciMETv3-cap experiment and the non-capture enzymatic conversion dataset without deploying any bias correction methodologies, producing comparable results for the distribution of cells in a reduced dimension representation, CH methylation distribution, and cell type composition between the experiments (Fig. 2B-E). Consistent with our previous sciMET-cap datasets, CG methylation was reduced compared to the genome-wide dataset due to the enrichment of regulatory regions that frequently exhibit hypomethylation and not due to conversion biases, which showed comparable global CH methylation levels (Fig. 2F).

Despite the increased fragment size using enzymatic conversion methods, we noticed a decrease in sequencing run quality with fewer clusters passing filter (<50% vs >90% typically). We suspected that this may be due to the unintentional conversion of sequencing adapter bases for the read 2 / index read 1 primer site that lies on the ligation junction between the indexed tagmentation oligo and indexed ligation oligo. To evaluate this, we performed Sanger sequencing using outer primers that are appended via PCR and are not subjected to conversion. This revealed distinct cytosine conversion to uracil at adapter bases present within the read 2 sequencing primer region (Fig. 2G). This is likely due to the sequence specificity of the TET2 catalytic domain, which biases its ability to protect methylated cytosines from conversion^12^. A possible solution to this problem would be the use of 5hmC (or other chemical modifications) in the adapter oligos to ensure protection; however, such modifications are costly and difficult to synthesize. Alternatively, the use of sequencing instruments that do not leverage this region for sequence read priming would eliminate the issue, such as a design compatible with the Ultima Genomics UG100^TM^ instrument.

### Atlas-scale dataset production is possible with sciMETv3 on multiple sequencing platforms

To demonstrate the atlas-scale potential of sciMETv3, we performed a single preparation on human brain specimens of four individuals (cortex, BA 46; 6596, 6926, 6996, 6998) which were distributed across equal numbers of tagmentation indexes (n = 24 each), providing the sample index in addition to the first tier of cell barcoding. After pooling, splitting and adapter ligation, and then pooling again, we obtained enough nuclei to dilute into 11 full 96-well plates at a target dilution count of 1,000 per well for an estimated potential cell count of just over 1 million. In total we diluted nuclei into six plates, four of which were banked for possible future processing (Fig. 3A). One plate was carried through bisulfite conversion, adapter ligation and PCR using primers established in previous experiments that append Illumina sequencing primers. The second plate was processed using bisulfite conversion for 88 wells, and 8 wells carried through enzymatic conversion. Adapter ligation was then performed followed by PCR using primers that append sequencing primers specific to the Ultima sequencing platform. Beyond alternate primer sequences, the other major design difference was to append the PCR index on the same side of the molecule as the tagmentation and ligation indexes so that the single-end reads produced by Ultima sequencing will read through all three indexes prior to the genomic DNA insert, maximizing the number of reads that will contain all three index sequences.

**Figure 3.**
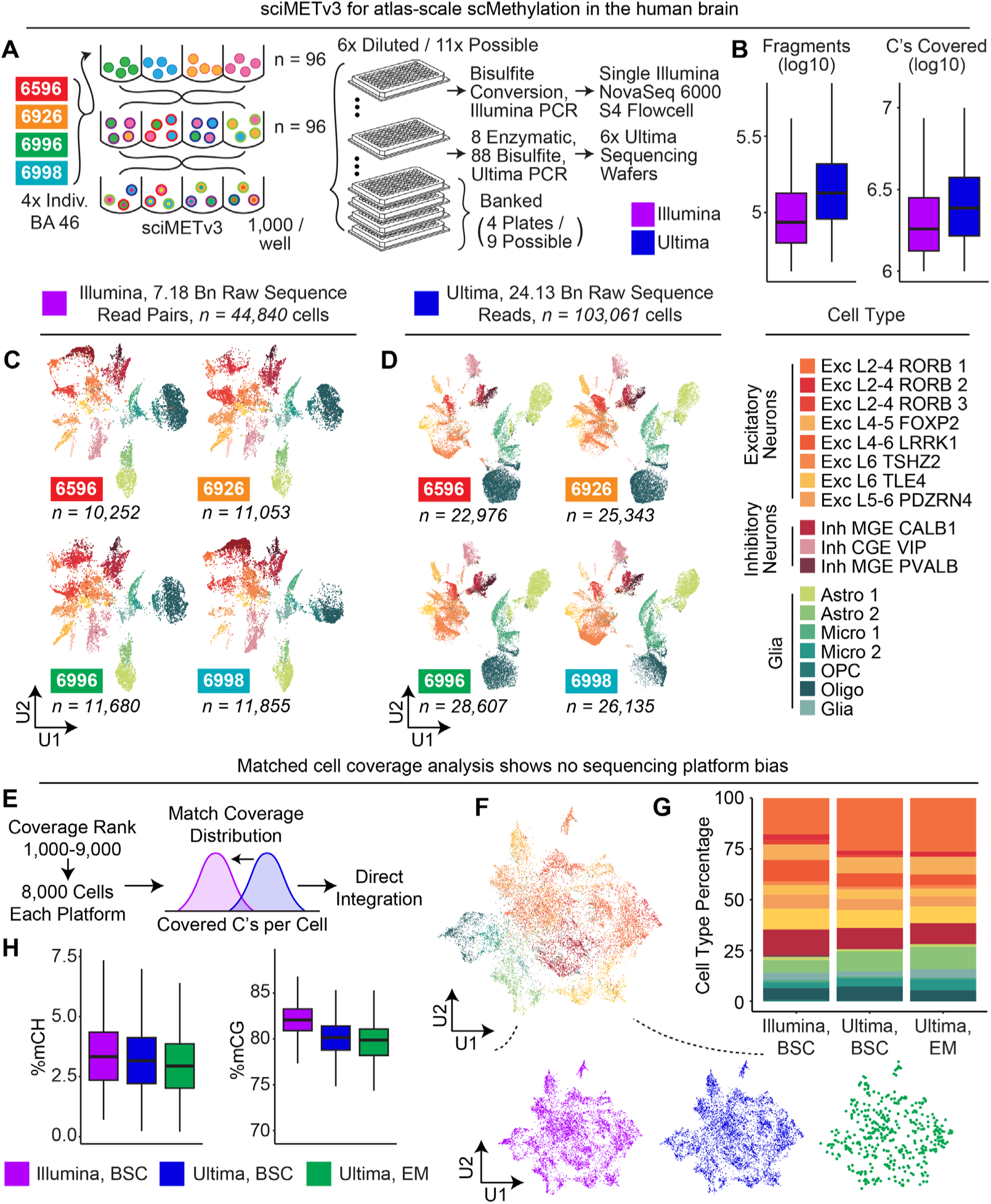
sciMETv3 can produce atlas-scale datasets using Illumina or Ultima sequencing platforms. **A)** Experimental design schematic. **B)** Summary of sequencing depth for each platform. **C)** UMAP of Illumina-sequenced cells split by four individuals and colored by cell type. **D)** UMAP of Ultima-sequenced cells. **E)** Strategy for matching cell coverage distribution between Illumina and Ultima sequenced cells for a direct comparison. **F)** UMAP of integrated coverage-matched cells from both platforms colored by cell type and split by platform (below). **G)** Comparable cell type proportions were achieved for each platform. **H)** Comparable global methylation statistics between platforms.

The first plate was sequenced on a single S4 flowcell of an Illumina NovaSeq 6000^TM^ instrument using a paired 200 cycle kit, producing 7.18 billion raw read pairs after demultiplexing. This resulted in 44,840 total cells called with a median of 1.82 million cytosines covered per cell at a median read duplicate rate of 13.98%, indicating that additional sequencing would yield greater coverage before reaching diminishing returns and increasing the total cell number with more cells reaching minimum coverage thresholds (Fig. 3B). Cells were split evenly across the four individuals (mean = 11,210 ± 6.4%) and clustering produced distinct primary cell types that were present in all individuals, consistent with previous observations that cell type specific methylation is the predominant signal that drives dimensionality reduction and clustering in brain single-cell DNA methylation datasets^7,13^ (Figs. 3C, S1).

The plate sequenced using the Ultima Genomics UG100^TM^ instrument was processed over six wafers, yielding a total of 28.5 billion raw reads, 24.13 billion after demultiplexing. The increased read counts over the Illumina-sequenced plate resulted in an increased median number of cytosines covered per cell, at 2.58 million with a commensurate increase in read duplicate rate, at 31.49%, producing 103,061 called cells (Fig. 3B). Similarly, cells were distributed evenly across all four individuals (mean = 25,765 ± 9.0%) with clustering driven by cell type over inter-individual variation (Fig. 3D). The lack of a need to preserve sequence integrity over the Illumina sequencing primer region using the Ultima platform enabled us to process a subset of the final indexing plate (n = 8 wells) using enzymatic conversion, which produced comparable coverage and methylation statistics when compared to the bisulfite converted cells (Fig. S2).

To evaluate any potential biases driven by the sequencing platform, we took the highest-covered 9,000 cells and then excluded the top 1,000 from each dataset, resulting in 8,000 cells for each platform. We then downsampled reads from the Ultima Genomics cells to achieve a matched distribution of cytosines covered per cell between each set (Fig. 3E). We then directly integrated the datasets without any batch correction methods, taking cells through windowing, dimensionality reduction and clustering, producing concordant distributions of cells across cell types for each platform, including enzymatic converted cells (Fig. 3F,G). We next assessed global methylation levels, which produced comparable CH methylation across both platforms and conversion methods, and a slightly reduced CG methylation level for both Ultima-sequenced conditions compared to the Illumina-sequenced cells (Fig. 3H). Taken together, sequencing platform and conversion method do not appear to produce any significant bias in the datasets.

### Integrated map of single-cell DNA methylation in the middle frontal gyrus from four individuals

We next leveraged all cells across both sequencing platforms to produce an integrated atlas of single- cell DNA methylation in the human middle frontal gyrus across four individuals, leveraging Harmony^14^ to account for the coverage differences between the two datasets (Fig. 4A,B). Clustering was performed followed by cell type assignment by correlation to a pre-existing atlas^13^ and assessing mCG patterns over canonical marker genes (Fig. 4C,D). The integrated atlas along with aggregated cell-type specific methylome profiles and all associated metadata is available as a downloadable R object for use as a reference map that enables interaction, visualization and integration using the Amethyst computational framework^15^.

**Figure 4.**
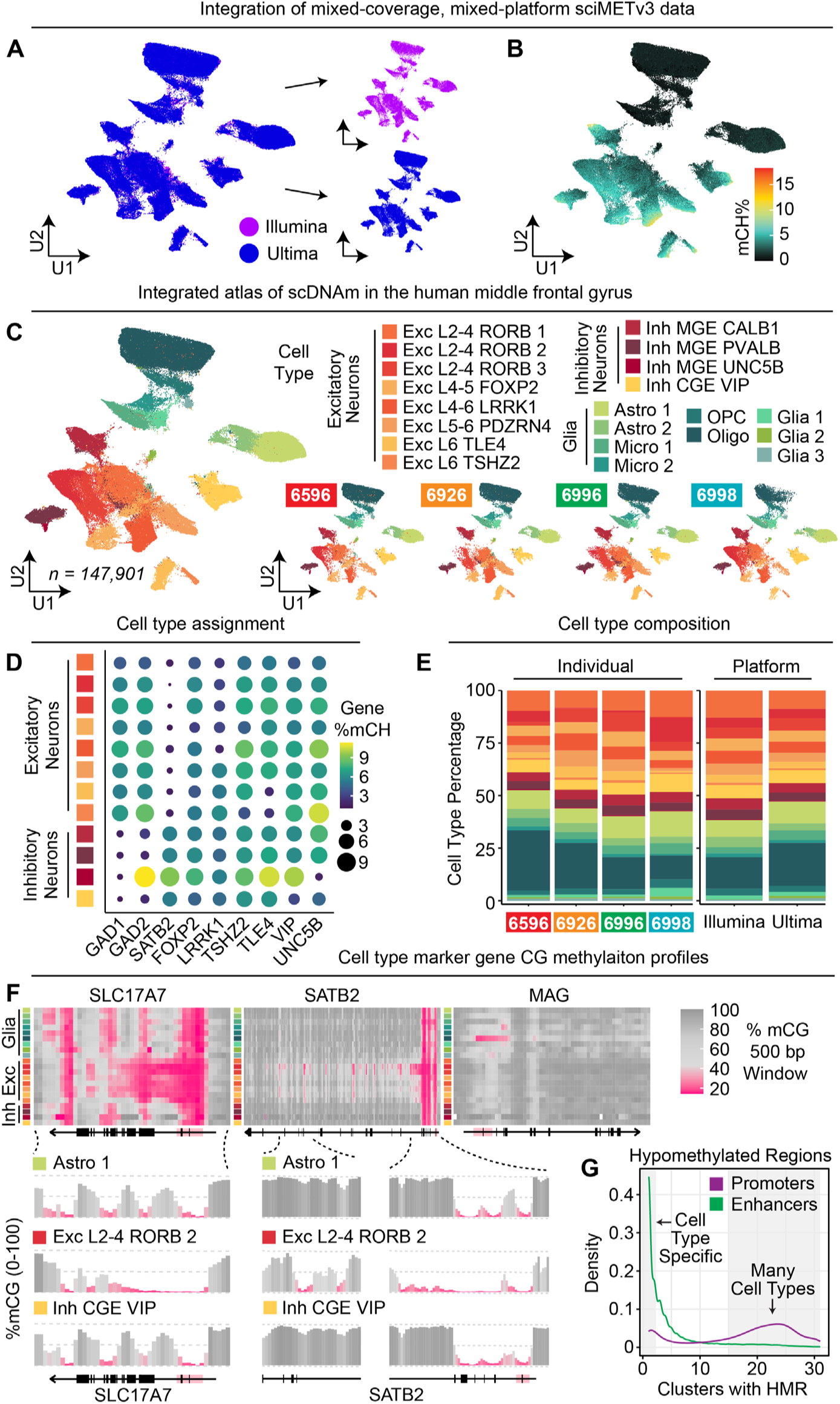
An atlas of single-cell DNA methylation in the human middle frontal gyrus. **A)** Combined UMAP across both sequencing platforms. **B)** Global mCH percentages for the combined dataset. **C)** Combined UMAP colored by cell type and split by individual (right). **D)** Marker gene body mCH levels by cluster. **E)** Cell type proportions across individual and sequencing platforms. **F)** mCG levels across marker genes show distinct cluster-specific patterns. **G)** Enhancers exhibit highly cell type-specific hypomethylation compared to promoters.

Cell type proportions were consistent across individuals as well as platforms, with the largest variance in the proportion of oligodendrocytes present (Fig. 4E). High-resolution aggregated CG methylation tracks were then generated for each cluster, providing a granular view of CG regulatory status genome-wide for each cell type. Similar to other epigenetic properties, such as ATAC-seq, DNA methylation status at promoters is varied across canonical marker genes, with some exhibiting cell type specific hypomethylation (e.g. MAG in oligodendrocytes), and others fully hypomethylated across all cell types. However, cell type- specific methylation patterning throughout the gene can be highly variable, with hypomethylation extending beyond the promoter and into the gene body, or in the form of focal dips in methylation throughout the gene (Fig. 4F).

To characterize these distinct patterns, we assessed cell type clusters (n = 31) genome-wide for hypomethylated regions (HMRs; methods). In total, 155,110 distinct HMRs were identified with 65,161 (42.0%) unique to a single cluster. Of these, 18,800 (12.1%) overlapped promoter regions with only 1,463 (7.8% of promoter HMRs) unique to a cluster and a mean of 17.5 clusters exhibiting hypomethylation at HMRs, indicating a propensity for cross-cell type promoter hypomethylation, regardless of expression status. In contrast, of the 44,304 (28.6%) of enhancer-overlapping HMRs, 14,668 (33.1% of enhancer HMRs) were cell type specific and a mean of 4.7 cell types exhibited hypomethylation at these HMRs, suggesting increased cell type specificity versus promoter elements (Fig. 4G).

### Step-wise indexed tagmentation enables DNA Methylation plus ATAC in single cells

We previously described a technology that enables the assessment of chromatin accessibility (ATAC) alongside whole genome sequence (WGS) from the same cells (scATAC+WGS) by leveraging two rounds of indexed tagmentation^16^. The first round of tagmentation is performed on native nuclei, thus capturing the open chromatin landscape. Subsequent fixation and nucleosome disruption enables the second round of tagmentation to be performed on the rest of the genome using a different index. Nuclei were then loaded onto a 10x Genomics Chromium instrument for droplet-based barcoding. Here, we applied a similar concept to our sciMETv3 workflow, performing an initial tagmentation on native human cortex nuclei using one set of 8 indexed sciMET Tn5 complexes. After the first round of tagmentation to encode open chromatin, we then performed fixation, nucleosome disruption and then a second round of tagmentation using a different set of 8 indexed complexes which are able to access the rest of the genome. Nuclei were then pooled and taken through the remainder of the sciMETv3 workflow, targeting 90 nuclei for each final well of indexing for an expected 6,480 cell profiles (Fig. 5A). Raw sequence reads were demultiplexed using the three tiers of indexing, splitting out the paired ATAC and MET indexes from the first round with 4.79% of reads derived from the first (ATAC) tagmentation and the remaining 95.21% from the second (MET) tagmentation, roughly matching the proportion of accessible versus inaccessible chromatin^17^. In total, 5,305 cells met minimum unique passing read counts for both the ATAC and MET paired datasets with read-depth concordance between the modalities (Fig. 5B).

**Figure 5.**
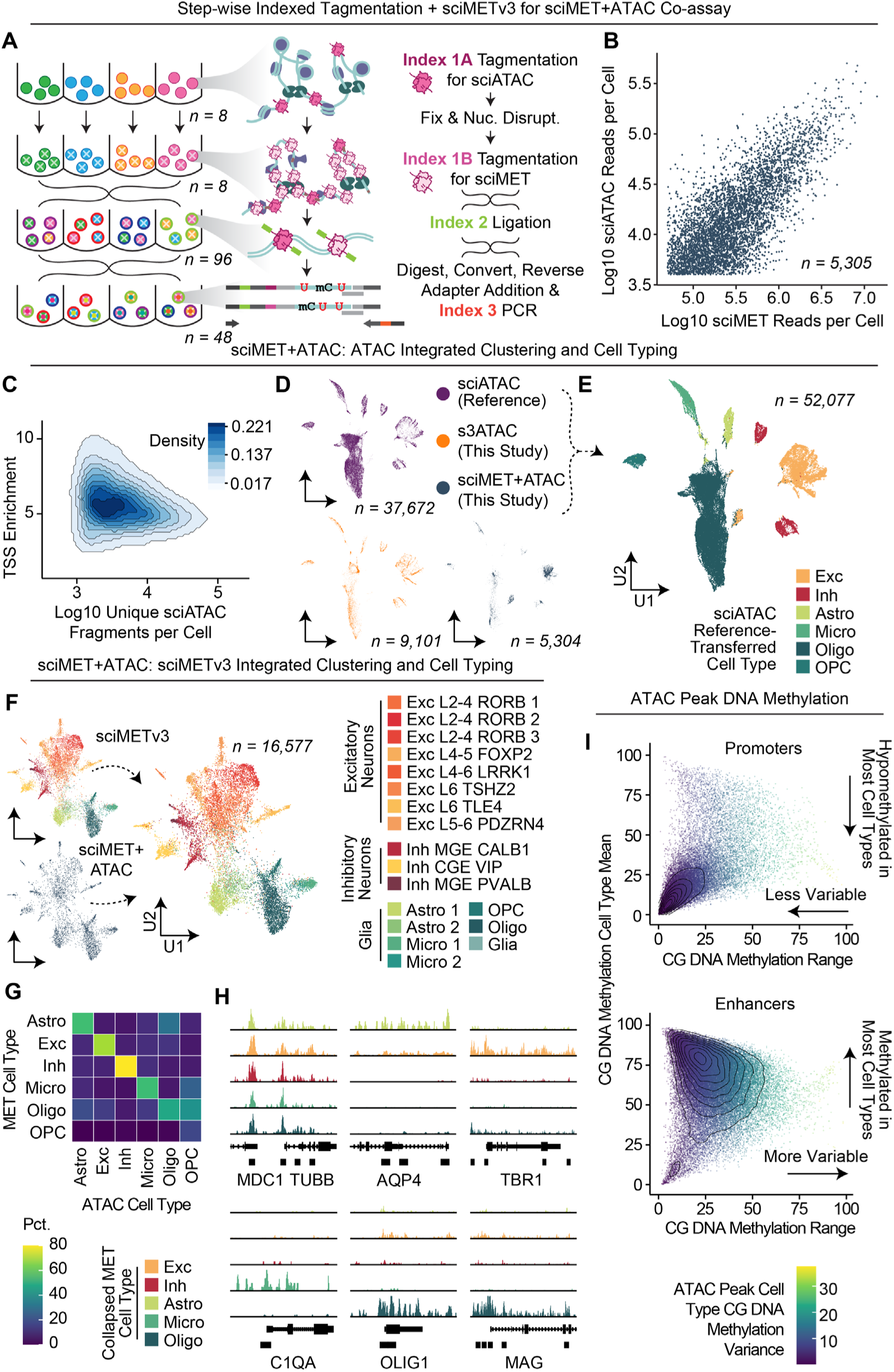
sciMET+ATAC for joint single-cell DNA methylation and chromatin accessibility. **A)** sciMET+ATAC co-assay schematic. **B)** Concordant ATAC and methylation read counts per cell. **C)** TSSe for the ATAC modality is low, yet consistent for the tissue sampled. **D)** UMAP of ATAC modality including a reference atlas and s3-ATAC preparation, split by dataset. **E)** ATAC-based UMAP colored by cell type. **F)** DNA methylation modality integrated with sciMETv3 reference cells from the same individual and colored by cell type. **G)** Cross- modality cell type concordance. **H)** ATAC profiles of marker genes split by DNA methylation-derived cell type. **I)** Called ATAC peaks at promoter regions exhibit less CG methylation variability between cell types versus putative enhancer peaks with higher cell type specificity.

ATAC reads were processed using the standard sciMET processing workflow through alignment. As an initial assessment, peaks were called using Macs2 ^18^, which produced 147,176 peaks from the 56.1 million total fragments, within the expected range for bulk ATAC-seq studies. Of these, 139,769 (95.00%) overlapped with previously identified accessible genomic loci, suggesting that the majority are likely *bona fide* candidate cis-regulatory elements^19^. Fragments were then used as input into SnapATAC2 ^20^ for single- cell level analysis. Transcription start site enrichment (TSSe) was relatively low (5.2; Fig. 5C) compared to typical single-cell ATAC-seq methods (∼10-20)^20^; however, this is expected due to the double tagmentation nature of the assay. For a typical scATAC workflow, two proximal tagmentation events are required in order to produce a short fragment that can be taken through subsequent library processing, with spurious tagmentation events yielding long fragments that are not able to be amplified in the final PCR stage. In our assay, spurious tagmentation events during the ATAC tagmentation are subjected to shortening due to the subsequent genome-wide tagmentation after nucleosome disruption, making them viable for downstream processing.

One valuable utilization of the sciMET+ATAC assay is the ability to leverage the ATAC modality for integration with existing reference atlas datasets where a methylation reference may not be available. We therefore generated an s3-ATAC^21^ dataset from the same tissue specimen using a novel implementation of the workflow that utilizes the iCell8 instrument for post-tagmentation processing in a 5,184 nanowell chip, similar to previous workflows for sciATAC^22^. In total, we leveraged a 32 × 32 nanowell setup targeting just under 12 pre-indexed nuclei per well for a total target of 12,000 total s3-ATAC profiles. Sequence reads were processed as above, producing 9,101 passing cell profiles with a relatively low TSSe of 5.6, suggesting tissue preservation may be a factor. We next leveraged the s3-ATAC profiles, the ATAC modality from the sciMET+ATAC assay, and an additional annotated reference dataset of ∼37 thousand cells, enriched for NeuN(-), (∼85%)^23^ to produce integrated clustering and visualization, using the annotations from the reference atlas to assign cell types to each cluster (Figs. 5D,E, S3A).

We next processed the DNA methylation side, producing cell groupings similar to the assigned cell types from the ATAC modality (Fig. 5F). The methylation modality was combined with our previous sciMETv3 dataset produced on the same individual, which produced substantial overlap except for a single cluster that was able to be filtered out using our doublet detection model, suggesting elevated noise in the dataset compared to the unimodal sciMETv3 workflow (Fig. S3B-D). We then leveraged the cluster identities from the unimodal dataset, as annotated in Figure 4, and performed label transfer to the sciMET+ATAC cells (Fig. 5F). Using the ATAC and MET cell type classes, we next compared cross-modality assignments which were largely concordant, including when leveraging the higher-granularity methylation-based clusters, with the exception of modest crosstalk between oligodendrocyte and oligodendrocyte precursor (OPC) cell populations (Fig. 5G).

Paired ATAC and genome-wide DNA methylation enables the assessment of both open and closed chromatin for DNA methylation status, as opposed to methods that conduct bisulfite conversion only on ATAC-derived reads, providing insight into the regulatory status of loci across all cell types and not just those that exhibit open chromatin. To assess these interactions, we leveraged the methylation-based cell typing to produce aggregated ATAC tracks, producing distinct cell type-specific accessibility patterns at marker genes (Fig. 5H). We then assessed ATAC peaks called from the data for methylation status across cell types, splitting out the ATAC peaks by promoters and enhancers (Fig. 5I). Between these categories, methylation was less variable at promoter regions, with nearly all cell types exhibiting hypomethylation. This low-variance hypomethylation population was present in the enhancer peak set, yet only for a minority of peaks, with the large majority exhibiting higher methylation variance where a majority of cell types exhibited hypermethylation.

## Discussion

Here, we describe sciMETv3, a robust technology for the production of atlas-scale single-cell DNA methylation datasets capable of delivering library sizes in the 100’s of thousands of cells. We demonstrate that sciMETv3 is compatible with capture-based techniques which allow for a reduced amount of sequencing to produce robust cell type clustering. Our assessment allowed for approximately 8,000 single-cell libraries to be multiplexed within a single capture reaction without a reduction in on-target capture rate. Notably, the capture workflows produce sufficient off-target coverage to provide genome-wide methylation calls when cells are aggregated at the cluster level, mitigating the limitation of capture techniques where non-targeted regions are missed.

The higher cell counts in the final indexing stage of sciMETv3 (∼600-1,000) over its predecessor, sciMETv2 (15-60), makes alternative means of C to T conversion viable, including EM-seq methods. We demonstrate the use of EM-seq on sciMETv3 libraries which produced a slightly larger fragment length which is likely due to the gentler treatment of the DNA by enzymatic steps versus the harsh chemical treatment with sodium bisulfite. Resulting libraries produced comparable methylation profiles and did not exhibit any bias in clustering and cell type proportions when compared to standard bisulfite-based conversion libraries. This result was confirmed by leveraging enzymatic conversion for libraries prepared using protocols for Ultima Genomics sequencing, where results were again indistinguishable from bisulfite-based converted libraries. However, we observed over-conversion of sequencing adapters which impeded Illumina sequencing which was not a factor using the Ultima platform due to the use of alternate primer regions.

We then demonstrate the production of a large-scale dataset produced from four human cortex specimens (middle frontal gyrus). Libraries were sequenced on either an Illumina NovaSeq 6000^TM^ or Ultima Genomics UG100^TM^ instrument with no discernable bias observed between the platforms. Notably, the single-end long read length nature of the UG100 instrument allows for minimal over-sequencing of internal bases within library fragments that get sequenced twice using paired-end sequencing where paired reads overlap. Achieving a longer fragment length could mitigate this observation, though even with enzymatic conversion methods a substantial number of fragments would exhibit overlapping coverage using the paired 200 bp sequencing format that we used in this study. Integration of all cells sequenced from this preparation yielded a high-resolution atlas of cell types in the human middle frontal gyrus, producing genome-wide maps of methylation profiles for each identified cell type.

Finally, we leverage a double-tagmentation workflow using two rounds of indexed Tn5 complexes with methylated adapters and an intervening nucleosome-disruption step. This workflow, sciMET+ATAC, enables the first tagmentation index to be leveraged for assessing chromatin accessibility, and the combination of both to be used as genome-wide DNA methylation. Overall, the data quality of sciMET+ATAC is lower for each modality than when performed on their own, as represented by a lower TSS enrichment value in the ATAC modality and the presence of noise in the methylation modality. However, the use of tailored quality control filtering allowed for distinct cell type identification, bolstered by integration with reference sciMETv3 cells from the same individual. Similarly, the ATAC modality integrated with an s3-ATAC dataset produced on the same tissue specimen using a novel nanowell chip-based implementation of the s3- ATAC workflow, as well as with an annotated reference dataset, which enabled cell type label transfer to the sciMET+ATAC cells. Notably, the DNA methylation modality was able to produce a higher granularity of neuronal clusters, likely due to the richness of CH methylation across the genome and the high information content produced using the sciMET assay. Taken together, we believe that the sciMET+ATAC workflow will be a valuable for profiling a portion of cells in addition to the sciMETv3 workflow to bridge between datasets and facilitate cross-modality integration and cell type assignment.

## Resource availability

All protocols are provided and all reagents are commercially available.

## Materials availability

All materials used in this study are readily available from commercial vendors.

## Author Contributions

R.V.N., B.L.O. and A.C.A. conceived the sciMETv3 and sciMET+ATAC technologies. R.V.N. performed all sciMETv3 and sciMET+ATAC preparations with assistance from B.L.O. B.L.O. performed the s3-ATAC preparation and developed the nanowell-based s3-ATAC workflow. S.N.A. performed capture experiments. L.E.R., B.L.O. and A.C.A. performed data analysis. Z.R. and N.I. performed Ultima Genomics sequencing and primary data processing of those data. A.C.A. supervised all aspects of the study and wrote the manuscript with input from all authors.

## Acknowledgements

Funding for this work was provided by NIH BRAIN Initiative RF1MH128842 and a Silver Family Foundation Innovator Award to A.C.A. Sequencing on the Ultima Genomics UG100^TM^ instrument was carried out at Ultima Genomics on libraries provided by the Adey Lab. We would like to thank Doron Lipson, Ph.D., Ariel Jaimovich, Ph.D., Alix Cruise, Ph.D. and Mirna Jarosz, Ph.D. of Ultima Genomics for facilitating the collaboration and providing sequencing data.

## Competing Interests

A.C.A. is an author of one or more patents that pertain to sciMET technology and an advisor to Scale Biosciences. This potential conflict is managed by the office of research integrity at OHSU.

## Data and Code Availability

All raw and processed data for this study are available in public repositories with unrestricted access. Raw data can be accessed from the NCBI Sequence Read Archive (SRA) under project accessions: PRJNA1126272 (sciMETv3) and PRJNA1134352 (sciMET+ATAC). Processed data can be accessed from the NCBI Gene Expression Omnibus (GEO) under accessions: GSE273592 (sciMETv3) and GSE272699 (sciMET+ATAC). All analysis was performed using publicly available software. DNA methylation analysis was carried out using Amethyst^15^ which is publicly available on GitHub: github.com/lrylaarsdam/amethyst.

## Methods

### Tissue homogenization and nuclei isolation

The brain tissue was dounce homogenized using cold NIB-Hepes buffer (10 mM Hepes, pH 7.5, 3 mM MgCl2, 10 mM NaCl, 0.1% IGEPAL (v/v), 0.1% Tween-20 (v/v), 1x protease inhibitor) as in Nichols et al. 2022. The cell suspension was then spun down (5 minutes, 500xg, 4C). The pellet was then resuspended in NIB-Hepes for nuclei quantification.

### Nucleosome disruption

Nuclei were quantified using a K2 Cellometer. Samples were separated into 1 million nuclei aliquots. Each aliquot was taken through the ScaleBio DNA Methylation Kit protocol for fixation and nucleosome disruption following manufacturer’s instructions. Afterwards nuclei were spun down at room temperature and resuspended in NIB-H. Aliquots were then recombined and quantified.

### Barcode 1: tagmentation

We tagmented 10,000-50,000 nuclei per well in a 96-well plate using Tn5 loaded with adapters containing all methylated cytosines (ScaleBio Part No: 941770). Each well contained 10 µL tagmentation buffer (ScaleBio Part No: 941788). The plate was incubated at 55°C for 15 minutes and then placed on ice. All wells were pooled and put into a 5 mL tube. 2 mL cold NIB-H was added, and the mixture was spun down at 500xG 4C for 5 minutes. The supernatant was removed. The mixture was washed with cold NIB-H + 3 μL BSA, spun down, and the supernatant was removed. The nuclei were then resuspended in 110 µL cold NIB-H, quantified, and used for in situ ligation.

### Barcode 2: ligation

To the 110 µL of nuclei, the following was added: 33 µL 10X Polynucleotide Kinase Buffer, 33 µL 10 mM ATP, 22 µL dH2O and 132 µL T4 Polynucleotide Kinase. The mixture was mixed by pipetting and distributed to a plate at 3 µL per well. The plate was incubated at 37°C for 30 minutes and then placed on ice. 2 µL of 15 µM ligation barcodes were added to each well of the plate. The following was then added to each well of the plate: 6.2 µL 2X StickTogether Buffer, 0.3 µL 100 µM v3 ligation splint and 1.5 µL T7 DNA Ligase. In other versions/experiments, the nuclei were kept in a 1.5 mL tube for the PNK 37°C incubation, after which the ligation master mix was added and the nuclei distributed to the plate containing the 96 ligation barcodes. The plate was incubated at 25°C for 1 hour and then placed on ice and allowed to cool fully. A full list of ligation oligo sequences can be found in Supplementary File 1.

### Post-ligation & dilution

All wells were pooled into a 5 mL tube. 3 mL NIB-H and 3 µL BSA were added. Nuclei were then spun down at 4°C 500xG for 5 minutes. The supernatant was removed. 3 mL NIB-H (with no protease inhibitors) was added. The tube was then spun down at 4°C 500xG for 5 minutes and resuspended in 100 µL NIB-H (no protease inhibitors). Nuclei were quantified and diluted to 750 nuclei per µL and 1 µL was added to each well of the final plates for bisulfite conversion using the ScaleBio Methylation Kit Met Bisulfite Conversion Module (Part No: 943631). Final plates or wells that used enzymatic conversion had 1 µL Qiagen Protease and 1 µL 90 mM Tris. The plates were spun down briefly and frozen at -20°C.

### Bisulfite conversion (BSC), cleanup, and adapter ligation

The plates for bisulfite conversion were defrosted and spun down briefly to collect the liquid to the bottom of the wells. Plates were then incubated at 50°C for 20 minutes to digest the nuclei and reverse cross-links. Bisulfite conversion, cleanup and reverse adapter ligation was carried out using manufacturers protocols for the ScaleBio Single-Cell DNA Methylation kit (ScaleBio Part No: 943631, 944302 and 944376).

### Enzymatic conversion, cleanup, and adapter ligation

Enzymatic conversion was carried out using the NEBNext Enzymatic Methyl-seq Conversion Module. Plates were spun down and then incubated at 55°C for 15 minutes and 72°C for 20 minutes to inactivate the Qiagen Protease. Afterwards, the manufacturer’s protocol was followed for enzymatic conversion. Final elution was done using 10 µL EB and then carried through the ScaleBio Single-cell DNA Methylation Kit (ScaleBio Part No: 944376) workflow for adapter ligation.

### Barcode 3: indexing PCR

The indexing PCR was performed with the following recipe for each well of a 96-well plate: 10 µL 5X VeraSeq GC Buffer, 2 µL 10 mM dNTPs, 1.5 µL VeraSeq ULtra Polymerase, 24 µL dH2O, 0.5 µL EvaGreen 100X and 1 µL 1 µM i7 Flow Cell primer for a total volume of 39 µL. 1 µL of barcoded i5 primers was added separately to each well. A full list of primers can be found in Supplementary File 1. The plate was mixed and placed on a qPCR with the following thermal conditions: 98°C initial denaturation for 30 seconds, 98°C for 30 seconds, 57°C annealing for 20 seconds, 72°C extension for 20 seconds, 72°C plate read for 10 seconds (these last 4 steps were cycled until exponential amplification was seen). After PCR, 10 µL of each well was pooled and the pool was column cleaned and SPRI cleaned with equal volume of product to SPRI beads. The resulting library was quantified using Qubit and TapeStation. Libraries were sequenced on an Illumina NextSeq 2000^TM^ or Illumina NovaSeq 6000^TM^.

### Ultima indexing PCR

For Ultima-compatible libraries, indexing PCR was carried out as above but substituting primers that ensure all indexes are on the same side of the molecule and that contain the Ultima Genomics outermost amplification and sequencing primers. A full list of primers can be found in Supplementary File 1. The final plate was pooled and sequenced on an Ultima Genomics UG100^TM^ instrument using six wafers.

### sciMETv3 capture

We pooled an 8-strip of sciMETv3 library in a volume of 16 µL of water. We performed capture with standard blockers and 300ng of library material. In a tube we combined 4 µL methylome panel (Twist Human Methylome Panel, Twist Bioscience, 105520), 8 µL Universal Blockers (also known as standard blockers, Twist Biosciences, 100578), 5 µL Blocker Solution (Twist Biosciences, 100578), 2 µL Methylation Enhancer (Twist Biosciences, 103557) and 1 µg of library in a volume of 7 µL in a 1.5 mL Eppendorf tube. Tubes were dried down on low heat in a speed-vac for 15’ and checked every 15’ for about an hour.

A thermal cycler was programmed as follows: 95°C hold / 95°C 5’ / 60°C hold (lid 85°C). 20 μl of 65°C. Fast Hybridization Mix (Twist Biosciences, 104180) was added to tubes with dried down panel, library and blockers. The mixture was solubilized for an additional five minutes before transferring to a 0.2 mL PCR tube. 30 µL of Hybridization Enhancer (Twist Biosciences, 104180) was added, the tube was pulse-spun and then transferred to the hot thermal cycler. The reaction was hybridized for 16 hrs to account for the large size of the methylome panel. Subsequent washing and PCR amplification was carried out according to manufacturer’s protocol, using a 63°C wash temperature.

### sciMETv3+ATAC

Nuclei were isolated in the same way as above. We tagmented 100,000-500,000 nuclei per well in an 8-strip using Tn5 loaded with adapters containing all methylated cytosines (ScaleBio Part No: 941770). Each well contained 10 µL tagmentation buffer (ScaleBio Part No: 941788). The 8-strip was incubated at 55°C for 10 minutes with 400 RPM shaking and then placed on ice. Each well was transferred to its own 1.5 mL tube where they were fixed and nucleosome disrupted using the same protocol as for the full sciMETv3 version. After nucleosome disruption it is important to remove all of the supernatant without disturbing the pellet. For the second tagmentation a new set of 8 barcodes was used and the nuclei were tagmented using the same recipe as above. They were also tagmented at 55°C for 10 minutes with 400 RPM shaking and then placed on ice. All wells were then pooled and carried through all post-tagmentation steps of the sciMETv3 protocol. Final plates had 90 nuclei diluted per well.

### Read processing

Raw sequence reads produced using Illumina instrumentation were carried through barcode demultiplexing using unidex (github.com/adeylab/unidex) to produce barcode-corrected read name paired fastq files. Reads were then taken through adapter trimming using ‘premethyst trim’ (github.com/adeylab/premethyst), which leverages Trim Galore. Sequence reads produced using the Ultima Genomics instrument were processed using the Ultima Genomics demultiplexing software to produce unaligned cram files containing the read with adapter bases trimmed and error-corrected indexes as a special field. These crams were then converted to fastq files with barcodes included within the read name for downstream compatibility.

### Alignment and methylation call extraction

Fastq files were aligned using the ‘premethyst align’ wrapper using default parameters which leverages BSBolt^24^. Aligned bam files were deduplicated using ‘premethyst rmdup’ and then methylation call files were generated using ‘premethyst extract’, including a lenient minimum read count threshold of 10,000 since cells are later filtered using more stringent parameters at subsequent analysis steps. Call files were then packaged into h5 calls files using ‘premethyst export’.

### DNA Methylation analysis using Amethyst

Cell metadata ‘cellInfo’ files produced from ‘premethyst extract’ along with methylation call h5 files were used to generate an Amethyst analysis object using amethyst (github.com/lrylaarsdam/amethyst)^15^ and then filtered to include cells meeting minimum cytosine coverage levels (1M for atlas dataset, 500k for other datasets). An hg38 reference annotation file was added for gene-level coordinates with the ‘makeRef()’ function. Site-level information in the h5 files were cataloged by chromosome using ‘indexChr’. Window methylation matrices were then generated with ‘makeWindows’, both for CG using metric = ‘score’ and CH using metric = ‘percent’. For the large-scale datasets produced using the Illumina NovaSeq 6000^TM^ and Ultima Genomics UG100^TM^ instruments, 100 kbp windows were leveraged, expanding to 200 kbp windows for all other smaller-scale datasets. We then estimated the number of IRLBA dimensions to calculate for the CG and CH contexts using ‘dimEstimate()’ followed by producing an IRLBA matrix using the specified number of recommended dimensions for each respective context using ‘runIrlba()’. Effects of coverage bias on the irlba matrix were mitigated with ‘regressCovBias()’. From the result, distinct groups were identified with the Rphenograph-based ‘runCluster()’ function and umap coordinates were projected using ‘runUmap()’. Cell type identification of the resulting clusters was performed based on the consensus of the following modalities: mCG patterns over canonical marker genes using amethyst vis ualization functions ‘histograM()’ and ‘heatMap()’; mCH levels over canonical marker genes using functions ‘dotM’ and ‘dimM’; and correlation of mCH levels over subtype-specific gene subset^26^ to a reference atlas produced by the Ecker Lab^13^. Integration of the Illumina and Ultima datasets was carried out using Harmony performed on the irlba matrix. Cell type annotation for the sciMET+ATAC dataset for the methylation modality was performed by integration of the sciMETv3 Illumina dataset for the same individual, leveraging the previously-annotated cell types to label- transfer to the sciMET+ATAC cells using the amethyst function ‘clusterLabelTransfer()’.

### s3-ATAC sample extraction and barcoded tagmentation

Frozen human brain tissue ID: 6996 was minced on dry ice and added to a Dounce homogenizer on ice along with cold 2 mL NIBH, containing fresh protease inhibitors. The tissue was homogenized with 7 strokes with the ‘A’ pestle, incubated for 10:00 on ice, then treated with 7 strokes with the ‘B’ pestle. The lysate was then filtered through 70 µm and 40 µm cell strainers (pluriSelect 43-50070 and 43-50030) and centrifuged at 500rcfr for 6:00 to remove extranuclear debris. The pellet was resuspended in 0.5 mL NIBH and counted on a Revvity K2 cellometer.

We performed tagmentation by adding 300 µL 4x TD Buffer and 12 µL 1M D-glucosamine (Sigma Aldrich), and an additional 388 µL NIBH to the nuclei for a final volume of 1.2 mL. We then distributed the nuclei into a 96-well PCR plate, at 10 µL per well, before adding 1.5 µL of barcoded tn5 (ScaleBio)^21^ and tagmenting at 55°C for 15 minutes. The plate was then transferred to ice and incubated 5:00 before pooling the nuclei into a 5 mL tube and adding 3mL NIBH. The nuclei were then centrifuged for 6:30 at 500 rcf and washed with 3 mL NIBH plus 3uL 100mg/ml BSA. After washing, the nuclei were resuspended in 100 µL, and counted on the K2 cellometer. Nuclei were diluted to 340 nuclei / µL for loading on the iCell8.

### s3-ATAC iCell8 loading protocol

For this sample, all additions were in a 36×36 well format. Volumes should be doubled if running the protocol in 72×72 well mode to account for the additional wells. LNA/SDS mix^21^ was distributed into a 350v iCell8 chip (TakaraBio) at 35 nL per well. The chip was blotted, capped with RC film (TakaraBio) and centrifuged 10:00 at 2500 rcf 4°C between every step unless otherwise specified. 50 nL diluted cells were then added, followed by incubation at 65°C for 10:00 and 72° for 10:00 (note: exact temperature settings for the modified BioRad T-1000 thermocycler included with the iCell8 were based on the conversion tables in Appendix G: Designing Thermocycler Programs, contained within the iCell8-CX User Manual).

Following nuclear lysis, we dispensed the Quench/Linear Extension mix^21^ at 50 nL per well. The chip was then placed in the BioRad TC-1000 for the Adapter Extension step. Adapter switching conditions were: initial extension of 72°C for 10 minutes, initial denaturation at 98°C for 30 seconds, then 10 cycles of 98°C for 10 seconds, 59°C for 20 seconds, and 72°C for 1 minute, followed by a 72°C final extension for 1 minute and cooling to 10°C hold.

35 nL each i7 TrueSeq and i5 Nextera barcoded PCR primers (15 µM) were added to each well, then 100 nL PCR Master-mix^21^ was added to each well (note: this can be added in 1x100 nL dispense for 36x36 mode, but must be added in 2, 50 nL dispense steps for 72x72 well mode). Final amplification conditions were: 98°C for 45 seconds, then 13 cycles of 98° for 15 seconds, 57°C for 30 seconds, and 72°C for 30 seconds, finishing with a 72° final extension for 5 minutes.

After extension, the PCR product was extracted from the iCell8 chip by centrifugation with the provided funnel from TakaraBio, and a 300µL aliquot was SPRI cleaned with a 1:1 sequential SPRI clean (sequentially adding 100 µL 3 times to the aliquot with 2 minutes binding time between additions to improve the size selection effect), before being eluted in 30ul and quantified with Qubit DNA fluorometer HS kit (Invitrogen) and Aglient Tapestation D1000.

The purified library was sequenced on a NextSeq 2000 P3 kit, with the following cycle numbers: Read 1: 89 bp, Index 1: 10 bp, Index 2: 10 bp, Read 2: 129 bp.

### s3-ATAC analysis

Sequencing data were demultiplexed with unidex and aligned with bwa mem^25^. The file was sorted by cell barcode, PCR duplicates were removed, and a custom python script was used to check the BAM file header for errors and add the cell barcode to the ‘CB’ BAM tag for each read to allow for faster ingest with SnapATAC2. A fragments file was created using SnapATAC2’s make_fragment_file function, and then an AnnData object was created with the import_data function with default parameters except for setting sorted_by_barcode to True. QC plots (fragment distribution and TSS enrichment) were generated as recommended in the SnapATAC2 documentation, and the dataset was filtered to remove cells with TSS- enrichment less than 5. Feature selection, dimensionality reduction, and clustering were all performed according to the SnapATAC documentation’s recommended settings.

For cell type assignment, we used the HGAP dataset^23^ and co-processed it with the human brain data described above, as well as with the ATAC data from the MET+ATAC coassay, using the same process described above, with the addition of removing batch effects with Harmony^14^. After clustering, the cell type information from the HGAP data was used to assign an implied cell types to the new datasets. As an additional validation, we use scanpy’s tracksplots function to plot the accessibility of various brain cell type marker genes across the different clusters. The results were concordant with the annotation lift-over.

---

**Figure S1.**
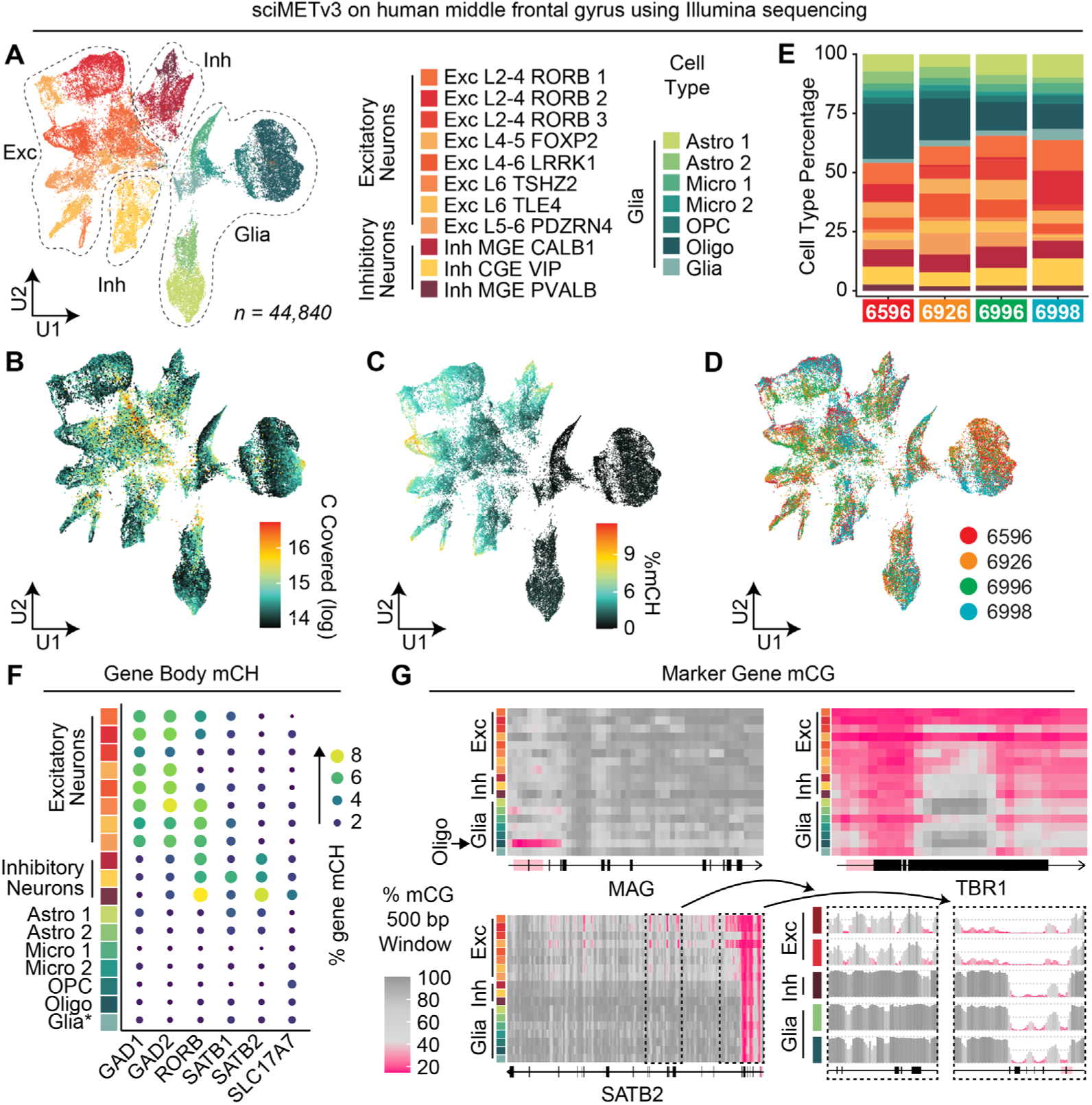
sciMETv3 sequenced on the Illumina platform. **A)** UMAP of cells colored by cell type. **B)** Cytosines covered per cell. **C)** Global mCH levels per cell. **D)** UMAP colored by individual. **E)** Cell type proportions by individual. **F)** Marker gene body mCH levels. **G)** Additional marker gene mCG methylation patterns with distinct cell type-specific hypomethylation regions.

**Figure S2.**
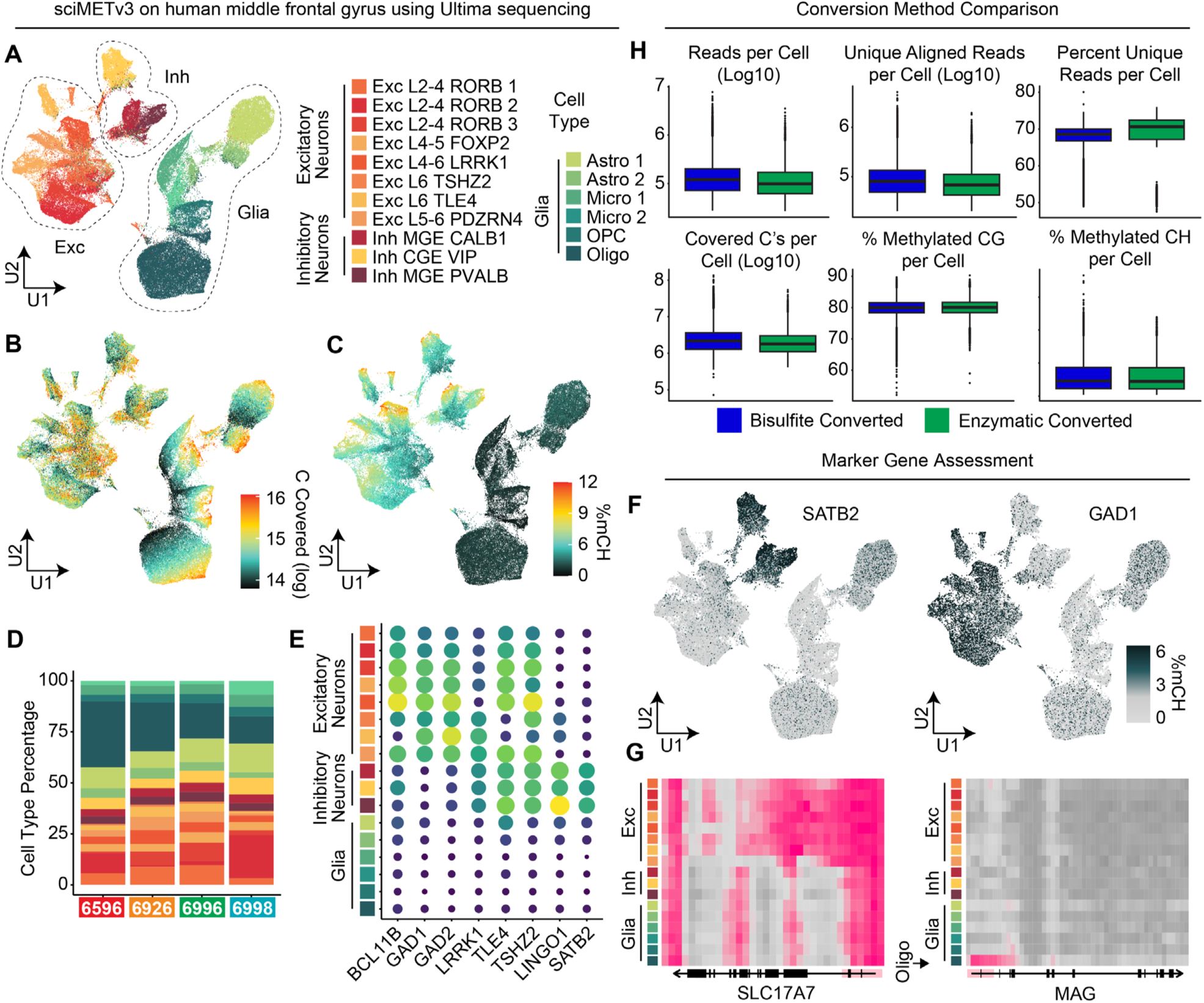
sciMETv3 sequenced on the Ultima platform. **A)** UMAP of cells colored by cell type. **B)** Cytosines covered per cell. **C)** Global mCH levels per cell. **D)** Cell type proportions by individual. **E)** Marker gene body mCH levels. **F)** mCH levels at the single-cell level projected onto the UMAP reveals inhibitory and excitatory neuron specificity. **G)** Additional marker gene mCG methylation patterns with distinct cell type-specific hypomethylation regions.

**Figure S3.**
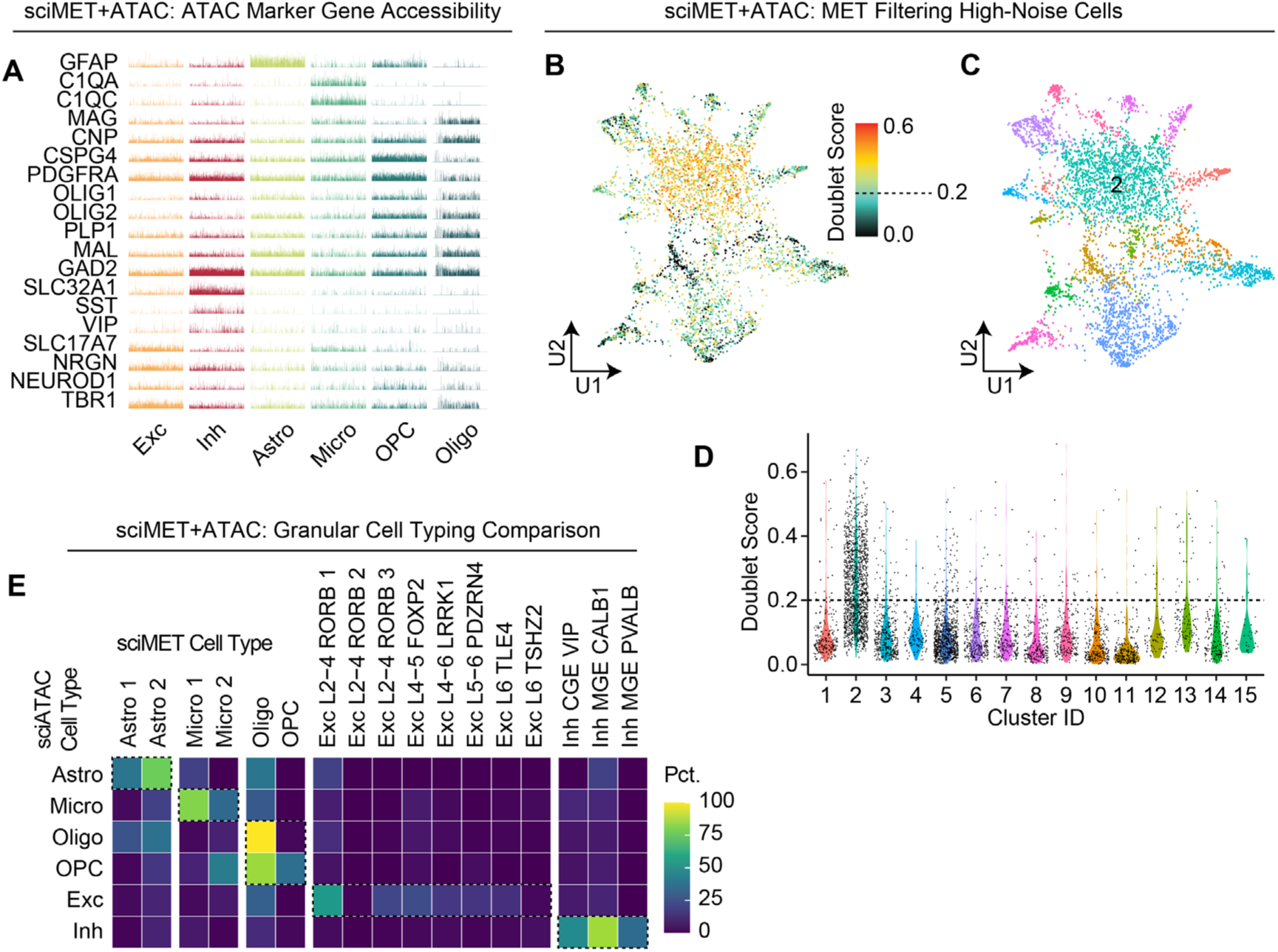
sciMET+ATAC cell typing and filtering. **A)** Marker gene tileplots for the ATAC modality. **B)** UMAP of sciMET+ATAC methylation cells reveals a population with a high doublet probability score. **C)** UMAP colored by cluster reveals that cluster 2 encompasses the high doublet score population. **D)** A score cutoff of 0.2 eliminates most of cluster 2 and other high-noise cells. **E)** Comparison of granular sciMET+ATAT DNA methylation-based cell types and ATAC-based cell types.

